# Analytical concordance of targeted next-generation sequencing and whole-genome sequencing for *Mycobacterium tuberculosis* drug resistance and lineage determination in West Java, Indonesia

**DOI:** 10.64898/2026.08.12.744373

**Authors:** Gusti Ayu Prani Pradani, Almira Alifia, Alia Putri Syahbaniati, Alamanda Larasmanah, Mohamad Busaeri, Hanif Djunaedy, Towifah Fauziah Choerunisa, Muhamad Nasrum Massi, Rifky Waluyajati Rachman, Azzania Fibriani, Reinout van Crevel, Jakko van Ingen, Bony Wiem Lestari

## Abstract

As drug-resistant tuberculosis (DR-TB) cases rise, resistance detection in a timely manner is essential to lead effective treatment and limit transmission. Targeted next-generation sequencing (tNGS) offers quick results with multiple important drugs covered, but assessments regarding its performance for DR-TB diagnostic use compared to whole genome sequencing (WGS) as the most comprehensive genomic-based tool are still limited.

This cross-sectional study compared resistance profiles generated by Deeplex Myc-TB tNGS assay with WGS for 116 prospectively-collected rifampicin resistant TB samples from West Java, Indonesia. All 116 samples were subject to paired analysis, the clinical samples were split to be directly processed for tNGS and to be cultivated for culture-based WGS. Both WGS and tNGS were carried out using Illumina MiSeq platform.

High concordance of tNGS and WGS were observed across thirteen anti-TB drugs evaluated, particularly for drugs included in the BPaLM regimen. Isoniazid had the lowest concordance of 86.73%. Of 116 samples, 31.03% (n = 36) had discrepant resistance calling from the two methods for one or more drugs, which came from 73 discordant variants identification. The most common source of discrepancy was when tNGS detected a resistance-conferring mutation while WGS did not (54.8%). tNGS could detect mixed infection better than WGS, but WGS was superior in identifying detailed major *Mycobacterium tuberculosis* lineage of the sample.

tNGS showed a good level concordance with WGS in detecting resistance-conferring mutations in rifampicin-resistant TB samples, with a more rapid turnaround time. Continuous update to tNGS panel and mutation catalogue is needed to keep the tool clinically relevant.

**Importance:** Drug-resistant tuberculosis (DR-TB) continues to pose worldwide threat, and newer diagnostic tools to generate quick, comprehensive resistance profile are crucial to provide timely appropriate treatment. Targeted next-generation sequencing (tNGS) is a promising new alternative, but more evidence on its performance is needed to support programmatic adoption. By analysing DR-TB samples with both tNGS and whole genome sequencing (WGS) and evaluating their results’ agreement, this study shows that tNGS works just as well as WGS in detecting TB drug resistance-conferring mutations, confirming its potential for routine diagnostic use. This study also observed that while WGS is superior in identifying *Mycobacterium tuberculosis* lineage with high resolution, it did not detect mixed infection better than tNGS. Notably, this study demonstrated that tNGS is clinically relevant for DR-TB detection in a high burden setting, providing evidence for programmatic consideration in Indonesia and other settings with similar demographics and TB situation.

## Introduction

Accurate identification of drug-resistance is essential in guiding drug-resistant tuberculosis (DR-TB) treatment. Inappropriately treated DR-TB cases would contribute to sustained high DR-TB prevalence in an area, increasing risk of wider DR-TB exposure and transmission (1). The threat of mistreatment and DR-TB primary acquisition are escalating as bedaquiline resistance is reported from multiple settings where BPaLM regimen is used (2). While molecular WHO-recommended rapid diagnostic (mWRD) tests such as Xpert MTB/RIF Ultra and Xpert MTB/XDR can detect drug-resistance much faster than phenotypic drug-susceptibility testing (pDST), their drug coverage is limited and does not cover most drugs in the BPaLM regimen (bedaquiline, pretomanid, linezolid, moxifloxacin) (3,4).

Genomic-based molecular diagnostics have recently transformed the possibility of rapidly identifying DR-TB. The whole genome sequencing (WGS) method enables the detection of a broad range of genetic variations across the entire *Mycobacterium tuberculosis* (Mtb) genome, providing a detailed genome-based resistance profile of the sample (5,6). Due to this comprehensive approach, WGS is considered standard for Mtb genetic and genomic studies (5). More recently, targeted next-generation sequencing (tNGS) method emerged as a simpler sequencing-based method alternative that can be used to detect DR-TB (7).

The simpler nature of tNGS compared to WGS has made it more promising for routine programmatic use. tNGS can process clinical samples directly without cultivation step, leading to substantially quicker turnaround time than WGS that requires high bacterial load from culture to provide interpretable results (7). Considering the potential of tNGS in TB control, the World Health Organisation (WHO) have included tNGS products of Deeplex Myc-TB, AmPORE-TB, and TBseq in their recommendations in the 2024 consolidated guidelines on tuberculosis for DR-TB testing, although it was stated that evidence level varies by drug and product (3). Currently, knowledge on how tNGS results compared to WGS beyond laboratory-confined accuracy studies are still limited.

In Indonesia, the national TB program has not yet incorporated sequencing technology to DR-TB testing algorithm (8). A number of studies have been done to assess feasibility of implementing tNGS for TB in the country, but insights regarding tNGS performance are needed to provide more evidence and would further aid clinicians in using and translating tNGS results in providing care. Therefore, this study aims to compare tNGS Deeplex Myc-TB results in identifying antituberculosis drug resistance with WGS as reference standard of genomic DST.

## Methods

### Study design and population

This is a prospective cross-sectional study comparing sequencing results of clinical samples obtained from tNGS and WGS methods, which was done as a sub-study to a larger implementation study of use of tNGS for DR-TB in West Java, Indonesia (9). All samples were rifampicin-resistance pulmonary TB confirmed by Xpert MTB/RIF Ultra assay. Samples were collected from hospitals and Community Health Centres (CHCs) in West Java, Indonesia from August 2024 to October 2025 as part of the West Java tNGS implementation study (9). All laboratory processes were done in West Java Provincial Health Laboratory, a provincial level referral laboratory for tuberculosis.

### Ethics

Participants were provided with verbal explanations and consenting participants gave written informed consent prior to sample collection. Ethical approval was obtained from the Health Research Ethics Committee at the Faculty of Medicine, Universitas Padjadjaran (97/UN6.KEP/EC/2025).

### Sample processing and sequencing

#### Specimen collection and pretreatment

Fresh clinical specimens (1–2 mL) were collected from eligible participants and decontaminated using standard N-acetyl-L-cysteine–sodium citrate–NaOH (NALC-NaOH) method (10). Decontaminated samples were resuspended in 3 mL phosphate-buffered saline, of which 1 mL aliquots were used for downstream testing: assessment of bacterial load and rifampicin resistance using the Xpert MTB/RIF Ultra assay (Cepheid, USA). DNA extraction for tNGS. and culture in Mycobacteria Growth Indicator Tube (MGIT; Becton Dickinson and Company, Franklin Lakes, NJ, USA). Confirmation test was performed on isolates grown on MGIT medium using the SD Bioline™ TB Ag MPT64 (Abbott Inc., South Korea) before cultivating on Lowenstein-Jensen (LJ) culture medium for subsequent WGS testing (10).

#### Targeted next-generation sequencing

DNA was extracted from sputum samples using a modified QIAamp DNA Mini Kit (Qiagen, Germany). The extracted DNA was then processed using Deeplex® Myc-TB assay (GenoScreen, Lille, France). After amplification, amplicon libraries were prepared with the Illumina DNA Prep Kit and sequenced on a MiSeq 100 plus platform (Illumina, San Diego, CA, USA). The Deeplex® Myc-TB pipeline was used for the analysis. The assay targets 18 main *Mycobacterium tuberculosis* gene targets associated with resistance to first-line drugs (rifampicin (RIF): rpoB; isoniazid (INH): inhA, gabG1, katG, ahpC; ethambutol (EMB): embB; and pyrazinamide (PZA): pncA), and second-line drugs (streptomycin (STM): gidB, rpsL, rrs; fluoroquinolones (FQ): gyrA, gyrB; kanamycin (KAN): rrs, eis; amikacin (AMK): rrs; capreomycin (CAP): rrs; ethionamide (ETH): inhA. fabG1, ethA; linezolid (LIN): rrl; bedaquiline (BDQ) and clofazimine (CFZ): rv0678).

#### Whole genome sequencing

Isolates grown in LJ medium between three and five weeks after the initial observation of growth. depending on visual assessment of colony density, were harvested using modified protocol to harvest all grown cultures from solid medium using 4 gr of sterilized glass beads (2 mm in diameter) in 4 ml of phosphate-buffered saline (PBS) (11).

DNA isolation was performed primarily according to the solid culture extraction protocol using InstaGene Matrix and high-speed Homogenizer, as demonstrated by Emilyn Conceição (11). Modification was made only to the bead beating step due to instrument availability. A BioSpec Mini-Beadbeater-16 Cell Disrupter was used with three cycles of 1 min at 3450 strokes per minute. with a one-minute interval between cycles. Additional purification was performed using a two-times AMPure ratio as described by Tomasz Suchan (11,12). The extracted DNA was then quantified using Qubit, and its purity was assessed using absorbance ratios (A260/A280 and A260/A230).

Sequencing libraries were prepared from purified DNA using Illumina DNA Prep Kit (13). Libraries were sequenced on a MiSeq platform using MiSeq reagent kit v2 (300-cycles). 25M with paired-end reads (2 x 150 bp). Sequencing was performed to achieve an average coverage depth of 50x (Illumina, San Diego, CA, USA).

#### Phenotypic drug-susceptibility testing

The phenotypic drug-susceptibility testing (pDST) was done as part of routine programmatic DR-TB management, and it is the current national standard for clinical decision-making and it was carried out following the national protocol from the Technical Guidance of Multiple Drug-Resistant Tuberculosis (MDR-TB) (8). Following this protocol, the pDST panel include bedaquiline (1 µg/ml), linezolid (1 µg/ml), clofazimine (1 µg/ml), isoniazid (0.1 µg/ml), pyrazinamide (100 µg/ml), levofloxacin (1 µg/ml), and moxifloxacin (0.25 µg/ml). However, during the study period, there were national situations that tentatively hinder testing of pyrazinamide. Additionally, a new regulation updated the pDST to be done for only bedaquiline, clofazimine, and linezolid from July 2025 onwards. These situations affect the availability of drug-specific pDST results of the samples included in this study.

### Data analysis

#### Sequencing depth

Sequencing parameters of both tNGS and WGS runs were obtained from the sequencers after each sequencing run was done. Sequencing depth was the main observed parameter which was used to determine whether the sequencing results were of adequate quality to be validly interpreted. Sequencing data from WGS run were rechecked using TBProfiler to ensure correct interpretation.

#### Drug resistance inference and discrepancies

Resistance profiles from tNGS were directly provided by the Deeplex Myc-TB software. while resistance profiles from WGS were processed using TBProfiler version 6.6.6. based on WHO TB catalogue and WHO TB mutation information sheets for genes carrying multiple resistance-associated mutations for different drugs (14,15). The depth of each gene in which the mutation was found was assessed using samtools version 1.21 (16).

The resistance profiles generated by both tNGS and WGS were then compared. Concordance was calculated for each drug by calculating the proportion of concordant results (called as resistant, susceptible, or uncertain significance of resistant on both methods) over the total number of samples.

Samples with ‘uncharacterised’ results were classified as ‘susceptible’ for concordance analysis. When a variant was reported by only one method, the result was considered discordant and noted as a discrepancy. Cohen’s kappa coefficient and its 95% confidence interval (CI) were calculated in R software version 4.5.2 using the cohen.kappa() function from the psych package (version 2.6.5) (17,18). Levels of agreement based on Cohen’s kappa would be ‘none’ (0—0.2), ‘minimal’ (0.21—0.39), weak (0.4—0.59), moderate (0.6—0.79), strong (0.8—0.9), and almost perfect (>0.9) (19). Prevalence-adjusted and bias-adjusted kappa (PABAK) was also calculated in R (20).

Positive percent agreement was calculated as the proportion of samples classified as resistant by tNGS out of samples classified as resistant by WGS. Negative percent agreement was calculated as the proportion of samples classified as susceptible by tNGS out of samples classified as susceptible by WGS. For positive and negative percent agreements, samples with ‘uncertain significance’ results were classified as ‘resistant’ as they are more likely to cause resistance although the degree of significance is not yet clear. For each measurement, corresponding 95% CI was estimated using the Wilson score method for binomial proportions via the binom package (version 1.1.2) in R (21).

#### Lineage identification

Lineage information of each sample processed by tNGS were provided by Deeplex Myc-TB software. Lineage identification from samples processed by WGS was obtained using analysis performed by TB-Profiler version 6.6.6 (14,15).

## Results

From 122 samples collected, two samples encountered failure in the WGS processes and could not generate results. Of the remaining 120 samples, four had their WGS sequencing depth lower than 30x, failing to meet the 30x threshold for the sequence reads to be deemed of good quality. Threfore, a total of 116 samples were considered qualified for paired tNGS-WGS analysis. The flowchartof sample inclusion to be analysed can be seen in Figure 1.

**Figure 1.**
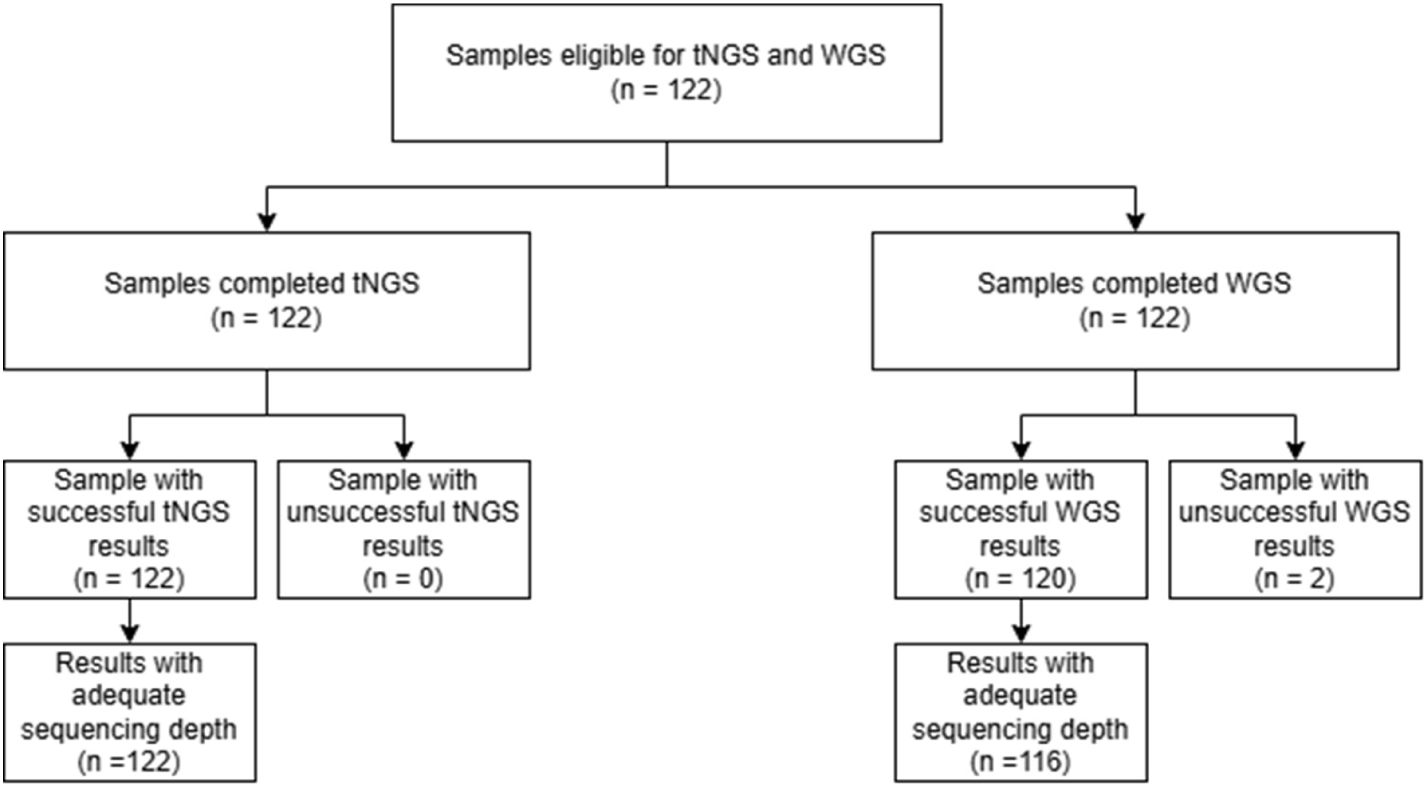
Sample eligibility flowchart.

### Sequencing depth

From the 116 samples, sequencing depth of WGS results were normally distributed with a mean and standard deviation of 75.50 ± 26.96 (IQR 56—95). Sequencing depth of results from tNGS were not normally distributed, with a median of 888.09 (IQR 489.21–1.220.45).

### Drug resistance inference and discrepancies

Concordance and Cohen’s kappa coefficients of tNGS results compared to WGS results are shown in Table 1.

**Table 1.** Analytical concordance of tNGS results in comparison with WGS results.

| Drug | n<br>paired | tNGS R |  |  | tNGS S |  |  | tNGS U |  |  | % Concordance<br>(95% CI) | Cohen's Kappa<br>(95% CI) | PABAK |
| --- | --- | --- | --- | --- | --- | --- | --- | --- | --- | --- | --- | --- | --- |
|  |  | R | S | U | R | S | U | R | S | U |  |  |  |
| WGS |  |  |  |  |  |  |  |  |  |  |  |  |  |
| Linezolid | 87 | 0 | 0 | 0 | 0 | 86 | 0 | 0 | 0 | 1 | 100 (95.8-100) | 1 (1 - 1) | 1 (0.92 – 1) |
| Amikacin | 108 | 1 | 0 | 0 | 0 | 106 | 0 | 0 | 1 | 0 | 99.07 (94.9-99.8) | 0.66 (0.05 - 1) | 0.98 (0.89 – 0.99) |
| Kanamycin | 108 | 1 | 0 | 0 | 0 | 104 | 0 | 0 | 3 | 0 | 97.22 (92.2 - 99.1) | 0.39 (-0.14 - 0.93) | 0.94 (0.84 – 0.98) |
| Fluoroquinolones | 115 | 14 | 4 | 0 | 0 | 97 | 0 | 0 | 0 | 0 | 96.52 (91.4 - 98.6) | 0.86 (0.72 - 0.99) | 0.93 (0.83 – 0.97) |
| Clofazimine | 112 | 2 | 3 | 0 | 0 | 104 | 0 | 0 | 1 | 2 | 96.43 (91.2 - 98.6) | 0.65 (0.34 - 0.97) | 0.93 (0.82 – 0.97) |
| Bedaquiline | 112 | 2 | 3 | 0 | 0 | 104 | 0 | 0 | 2 | 1 | 95.54 (89.9 - 98.1) | 0.53 (0.18 - 0.89) | 0.91 (0.79 – 0.96) |
| Pyrazinamide | 112 | 15 | 3 | 0 | 1 | 90 | 1 | 0 | 0 | 2 | 95.54 (89.9 - 98.1) | 0.85 (0.71 - 0.98) | 0.91 (0.79 – 0.96) |
| Ethionamide | 105 | 14 | 1 | 0 | 1 | 80 | 0 | 0 | 3 | 6 | 95.24 (89.3 - 97.9) | 0.87 (0.75 - 0.98) | 0.90 (0.79 – 0.96) |
| Ethambutol | 116 | 26 | 3 | 0 | 1 | 82 | 1 | 0 | 1 | 2 | 94.83 (89.2 - 97.6) | 0.87 (0.77 - 0.97) | 0.90 (0.78 – 0.95) |
| Capreomycin | 105 | 1 | 4 | 0 | 0 | 96 | 0 | 0 | 3 | 1 | 93.33 (86.8 - 96.7) | 0.35 (-0.01 - 0.71) | 0.87 (0.74 – 0.94) |
| Rifampicin | 116 | 105 | 7 | 0 | 1 | 2 | 0 | 0 | 0 | 1 | 93.1 (86.9 - 96.5) | 0.40 (0.07 - 0.73) | 0.86 (0.74 – 0.93) |
| Streptomycin | 110 | 22 | 3 | 0 | 2 | 75 | 0 | 0 | 4 | 4 | 91.82 (85.2 - 95.6) | 0.81 (0.69 - 0.93) | 0.84 (0.70 – 0.91) |
| Isoniazid | 113 | 60 | 8 | 0 | 2 | 35 | 2 | 1 | 2 | 3 | 86.73 (79.3 - 91.8) | 0.75 (0.63 - 0.86) | 0.73 (0.56 – 0.84) |

R = resistant. S = susceptible. and U = uncertain significance. Samples with 'uncharacterised' results were classified as susceptible.

Concordance between targeted NGS and WGS was generally high across all anti-tuberculosis drugs. ranging from 86.73% to 100% (Table 1). Highest concordance was observed in linezolid, while the lowest was isoniazid. Most concordant results consisted of susceptible-susceptible classifications. followed by tNGS resistant—WGS resistant, whereas tNGS uncertain significance—WGS uncertain significance classifications were relatively uncommon across all drugs. Despite the overall high concordance, out of 116 analysed samples, 37 (31.89%) samples had discrepancies in calling the resistance status of one or more drugs.

Based on the Cohen’s kappa coefficient, most drugs had a ‘strong’ level of agreement between tNGS and WGS (41.67%; 5/12), and linezolid had ‘almost perfect’ level of agreement with a note that no linezolid-resistant sample was detected by either tNGS or WGS. Two second-line injectable drugs were put in the ‘minimal’ agreement category while rifampicin and bedaquiline were categorised in the ‘weak’ level of agreement. However, these drugs had higher agreement levels when PABAK was used instead of Cohen’s kappa. PABAK coefficient were notably high across all drugs, except for isoniazid.

**Table 2.**
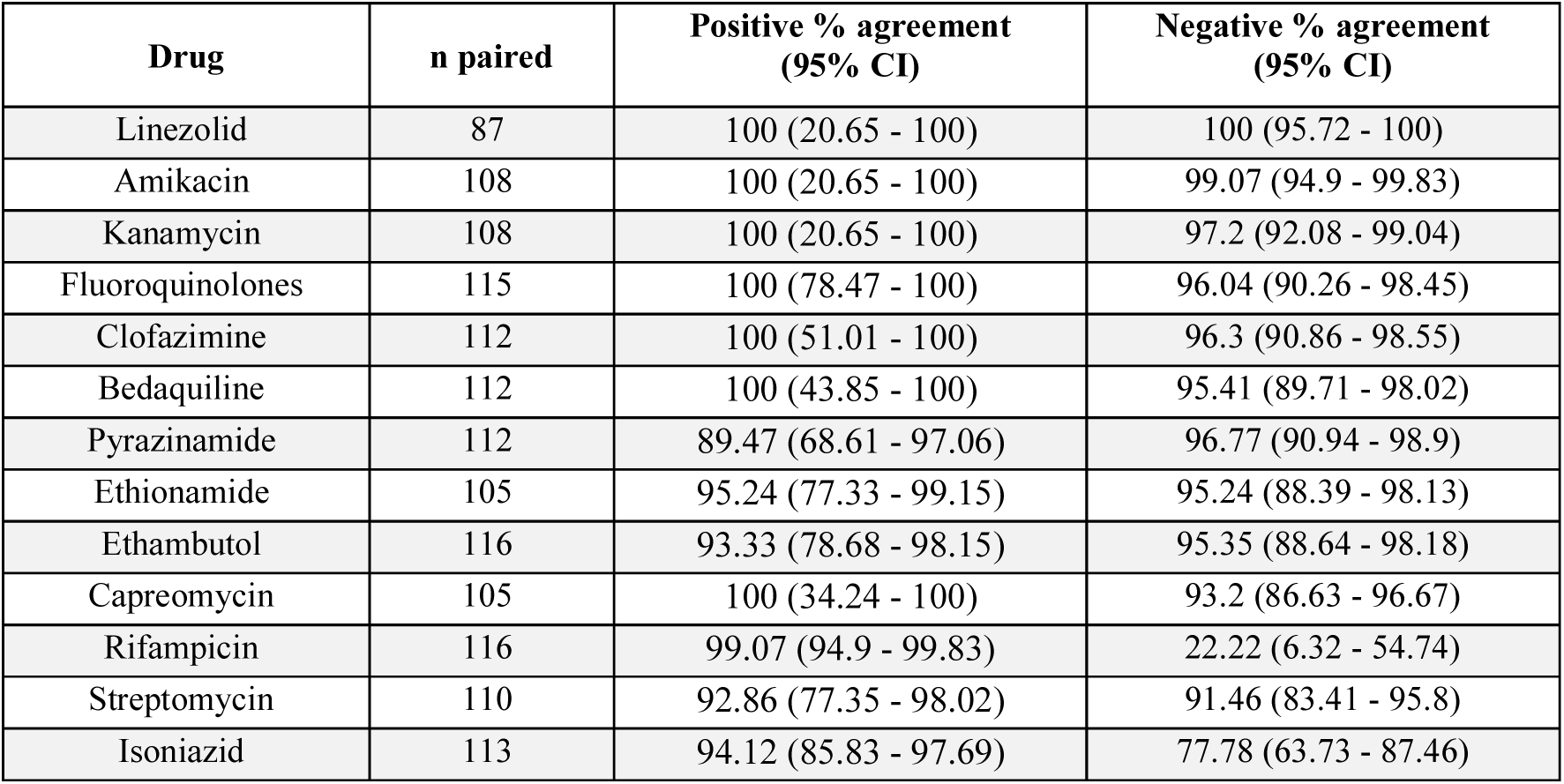
Positive and negative agreements of tNGS results and WGS results. Samples with ‘uncharacterised’ results were classified as susceptible, while ‘uncertain significance’ results were considered as ‘resistant’.

| Drug | n paired | Positive % agreement (95% CI) | Negative % agreement (95% CI) |
| --- | --- | --- | --- |
| Linezolid | 87 | 100 (20.65 - 100) | 100 (95.72 - 100) |
| Amikacin | 108 | 100 (20.65 - 100) | 99.07 (94.9 - 99.83) |
| Kanamycin | 108 | 100 (20.65 - 100) | 97.2 (92.08 - 99.04) |
| Fluoroquinolones | 115 | 100 (78.47 - 100) | 96.04 (90.26 - 98.45) |
| Clofazimine | 112 | 100 (51.01 - 100) | 96.3 (90.86 - 98.55) |
| Bedaquiline | 112 | 100 (43.85 - 100) | 95.41 (89.71 - 98.02) |
| Pyrazinamide | 112 | 89.47 (68.61 - 97.06) | 96.77 (90.94 - 98.9) |
| Ethionamide | 105 | 95.24 (77.33 - 99.15) | 95.24 (88.39 - 98.13) |
| Ethambutol | 116 | 93.33 (78.68 - 98.15) | 95.35 (88.64 - 98.18) |
| Capreomycin | 105 | 100 (34.24 - 100) | 93.2 (86.63 - 96.67) |
| Rifampicin | 116 | 99.07 (94.9 - 99.83) | 22.22 (6.32 - 54.74) |
| Streptomycin | 110 | 92.86 (77.35 - 98.02) | 91.46 (83.41 - 95.8) |
| Isoniazid | 113 | 94.12 (85.83 - 97.69) | 77.78 (63.73 - 87.46) |

Positive percent agreements were relatively high across all drugs, of around 90% and higher. Six drugs had perfect (100%) positive percent agreement, although wide CIs were noted especially for linezolid, amikacin, and kanamycin. Negative percent agreements were predominantly high (>90%) except for isoniazid and rifampicin.

Seventy-two discrepancy occurrences of resistance interpretation among tNGS and WGS affected twelve drugs (Table 3), with the most common discrepancy was tNGS resistant—WGS susceptible (RS) (54.2%, 39/72), and isoniazid (INH) was the most impacted drug by discrepancies with 15 occurrences, followed by rifampicin (n=8). Detailed information of each discrepancy can be seen in Supplementary Material 1.

**Table 3.** Summary of discrepancy results between tNGS and WGS. Note that Fluroquinolones here include either Levofloxacin or Moxifloxacin.

| Type of discrepancy | Total instances | Affected drugs | Average tNGS coverage depth | Average WGS coverage depth |
| --- | --- | --- | --- | --- |
| tNGS Resistant, WGS Susceptible | 39 | BDQ, CAP, CFZ, EMB, ETH, FQ, INH, PZA, RIF, STM | 836.27 | 80 |
| tNGS Uncertain significance, WGS Susceptible | 20 | AMK, BDQ, CAP, CFZ, ETH, INH, KAN, STM | 634.07 | 76.70 |
| tNGS Susceptible, WGS Resistant | 8 | EMB, ETH, INH, PZA, RIF, STM | 996.8 | 46.86 |
| tNGS Susceptible, WGS Uncertain significance | 4 | EMB, INH, PZA | 672.4 | 76.4 |
| tNGS Uncertain significance, WGS Resistant | 1 | INH | 422.58 | 74.25 |

Upon closer look at gene level, the 72 drug-level discrepancies came from 73 discordant resistance interpretations across 17 resistance-associated genes. The genes contributing the highest number of discordant interpretations were katG (11/73, 15.06%), rrs (9/73, 12.3%), rv0678 (9/73, 12.33%), rpoB (8/73, 10.96%), and embB (6/73, 8.22%). Together, these five genes accounted for approximately 62% of all discordant gene-level classifications. Most discordances in katG, rpoB, rv0678, and embB were classified as tNGS resistant while WGS classified as susceptible. In contrast, discordances in rrs, ethA, and gidB were predominantly driven by uncertain calls in WGS or tNGS. Notably, katG exhibited discordances across all five classification categories, indicating greater variability in resistance interpretation compared with the other genes.

Among the discordant results, the most frequently observed mutation was katG S315T (7 instances), Other recurring mutations included rrs g637a (4 instances), rrs c845t (3 instances), gyrA D94G (3 instances), and rpoB S450L (3 instances). Two instances of rv0678 Y157* were observed affecting bedaquiline resistance interpretations.

Of the 116 samples included in the genomic analysis, 26 lacked corresponding phenotypic drug susceptibility testing (pDST) results due to insufficient sample material for the test. Of the 90 samples that had available pDST results, almost half (45.56%, 41/90) were tested for only bedaquiline, linezolid, and clofazimine pDST panel. Of all available pDST data, both genomic approaches demonstrated high concordance with pDST for bedaquiline, linezolid, and clofazimine. Lower concordance was observed for isoniazid and pyrazinamide. The concordance values with pDST were higher for WGS compared to tNGS across most drugs (Supplementary Material 2). Among samples with discordant WGS and tNGS results, only 9 had their pDST results available and for only a limited number of drugs. Of those, WGS had more concordant results with pDST compared to tNGS (Supplementary Material 3).

### Lineage identification

Of the 116 paired samples, 115 had their lineage successfully identified by both WGS and tNGS, as tNGS failed to identify the lineage of one sample. Out of 115 samples, 73.91% (85/115) had their detected main lineage from tNGS in concordance with those of WGS. Of these concordant samples, 57 (67.06%) belonged to Lineage 2. predominantly sublineage 2.2.1. 27 (31.76%) belonged to Lineage 4. and one (1.18%) belonged to Lineage 1. Among samples with concordant main lineages. mixed-lineage infections were identified in 20 samples by tNGS and in six samples by WGS.

Lineage discordance was observed in 26.09% of the samples (30/115). The discordances mostly came from samples classified by tNGS as ‘Lineage 2’ while they were classified as ‘Lineage 4’ by WGS (n=14). Following that, 11 samples were specified as ‘Other than lineage 4.9’ by tNGS when those samples were identified as ‘Lineage 2’ by WGS.

## Discussion

This study evaluated multiple measurement of agreements between tNGS and WGS in detecting anti-TB drug resistance and found that tNGS demonstrated high concordance across most drugs, including for BPaLM regimen drugs which is currently the default for DR-TB treatment in Indonesia, despite the presence of minor discrepancies. These findings showed that tNGS performs comparably to WGS and has the potential to serve as a reliable diagnostic tool for the detection of drug-resistant tuberculosis.

The overall findings on the tNGS and WGS concordance are largely consistent with those reported by another study from South Africa, which reported an overall concordance of more than 92% across all drugs except isoniazid and ethionamide (4). Isoniazid also had the lowest concordance in this study, but a noticeably higher concordance of tNGS and WGS on ethionamide resistance calling was observed in this study which may be attributable to the larger number of ethionamide-resistant samples included in our samples. Linezolid was found to have the highest concordance in both studies—100% in this study and 97% in their study—the difference might come from potential overestimate in this study as there were no linezolid-resistant samples present in our cohort.

Using other means of measurements, the levels of agreement between tNGS and WGS were relatively high across all drugs. Kanamycin and capreomycin were noted to have low tNGS vs WGS agreements based on their kappa coefficient. This is quite expected, as predicting second-line injectable drugs susceptibility have been reported in multiple studies especially when it involves ribosomal genes like *rrs*, and the differences in pre-sequencing sample processing between tNGS and WGS can contribute to more observed discrepancy (22,23). Rifampicin had very low negative percent agreement in this study, which might came from the very low number of tNGS Susceptible—WGS Susceptible found in this study, as we methodologically only evaluated samples with confirmed rifampicin-resistance on Xpert MTB/RIF Ultra. The highly disproportionate numbers among resistance categories for rifampicin might also contributed to the drug’s observed low Cohen’s kappa, although when the kappa is adjusted for prevalence and systematic biases using PABAK, the corrected level of agreement seemed to reflect the observed findings better.

This study found that tNGS can generate better yields in term of successful sequencing results compared to WGS, and while several types of discrepancies were present, WGS contributed to the majority of missed mutations detection. Overall, WGS showed a much less sample coverage depth compared to tNGS, which is reasonable considering its mechanism to capture the whole genome. This could explain why the majority of discordant results came from the samples where tNGS captured resistance-associated mutations while WGS did not, as tNGS could sequence the targeted genes in more depth compared to WGS. However, there were several instances when the opposite happens, i.e. tNGS missed a resistance-associated mutation that WGS captured. This can be due to the isolate cultivation process done prior to WGS, which could promote growth of certain mutants and eventually alter the dominant population in the sample, making the final result differ from tNGS (23).

Based on this study’s findings, tNGS can be considered to be sufficiently reliable for diagnostic use. The high concordance with WGS showed that its performance is not inferior compared to WGS and despite the presence of discrepancies, WGS missed resistance-associated mutations in the tNGS-targeted area approximately 4-fold more often than tNGS. This situation is similar to other study from China that used tNGS assay from Shengshizhongfang (Beijing) Bio Sci & Tech. Co., Ltd., where they found the ratio of tNGS-only detected mutations versus WGS-only detected mutations to be around 2:1 (23). While this study observed WGS had more agreement with pDST results, it can be attributable to the cultivation process that was part of these two tests, and it is important to note that the number of available paired pDST results in this study were low and for a very narrowed selection of drugs only.

The targeted aspect of tNGS also introduced weakness as some of the resistance-associated mutations beyond the targeted area would not be covered. The most clinically relevant one being related to BPaLM regimen is mutations related to pretomanid, which is currently not targeted by Deeplex Myc-TB kit. However, WGS did not detect any pretomanid-resistant samples in this study, and most of WGS-only detected variants beyond tNGS target regions were also not significantly related to resistance and hence have minimal clinical impact. This aligns with another study from South Africa, that analysed more than 2,000 variants only reported by WGS beyond Deeplex Myc-TB targeted genes to be classified as non-resistance-conferring (4). However, while tNGS can currently be considered sufficient for diagnostic use, prevalence of pretomanid resistance and potentially other new resistance-conferring mutations may rise, noting that timely routine update to the kit and characterisation of more mutations are warranted to keep the clinical relevance and reliability of tNGS results.

In lineage analysis, we found that WGS provided greater phylogenetic resolution by assigning isolates to sublineage and sub-sublineage levels. In contrast, tNGS identified mixed-lineage populations more frequently than WGS, suggesting a greater sensitivity for detecting mixed infections. Both of these findings are consistent with several previous studies (24,25). Discordant lineage classifications between both techniques demonstrated potential limitations of tNGS in identifying the lineage and that it could report unclear results, which aligns with other studies (4,24). The discrepancy is reasonable considering WGS interrogates genome-wide SNPs, while tNGS relies only on a limited set of targeted loci. Therefore, WGS-based SNP classification is likely to provide a more robust phylogenetic assignment than spoligotype-informed tNGS calls and remains preferable when accurate discrimination between major lineages and sublineages is required for molecular epidemiology and transmission analysis. This advantage of WGS has been consistently demonstrated in previous studies comparing WGS with conventional genotyping methods (26–29). Nevertheless, tNGS remains useful for broad lineage screening and routine clinical use, when detailed lineage information is not vitally needed.

This is one of the first studies in Indonesia comparing tNGS and WGS using real programmatic samples of DR-TB patients in West Java, using quite a considerable number of samples for this study design.

This study has several limitations. First, this study is limited to West Java, Indonesia, which may corelate with the prevalence of mutations detected and informed in this analysis. Other regions might have certain mutations with different prevalence that might fall within or beyond the tNGS targeted sequencing region, affecting the generalisability of tNGS value in other settings. Second, this study is limited to the use of Deeplex Myc-TB kit assay for tNGS and TBProfiler for WGS analysis. Interpretations and comparisons with other studies using other platform and software must be done carefully considering potential interplatform differences in diagnostic performance and variant threshold calling.

In conclusion, this study demonstrates that tNGS performed well in detecting tuberculosis drugs-related mutations in comparison to WGS, showing its reliability and potential for programmatic diagnostic and clinical use, particularly with its simpler and more rapid processes compared to WGS.

## Data availability statement

Deidentified individual data of observed resistance-associated mutations and resistance status which were used as the basis of analysis of this study are available in the Open Science Framework (OSF) repository at https://doi.org/10.17605/OSF.IO/MF6VJ.

## Supplementary Materials

**Supplementary Material 1.**
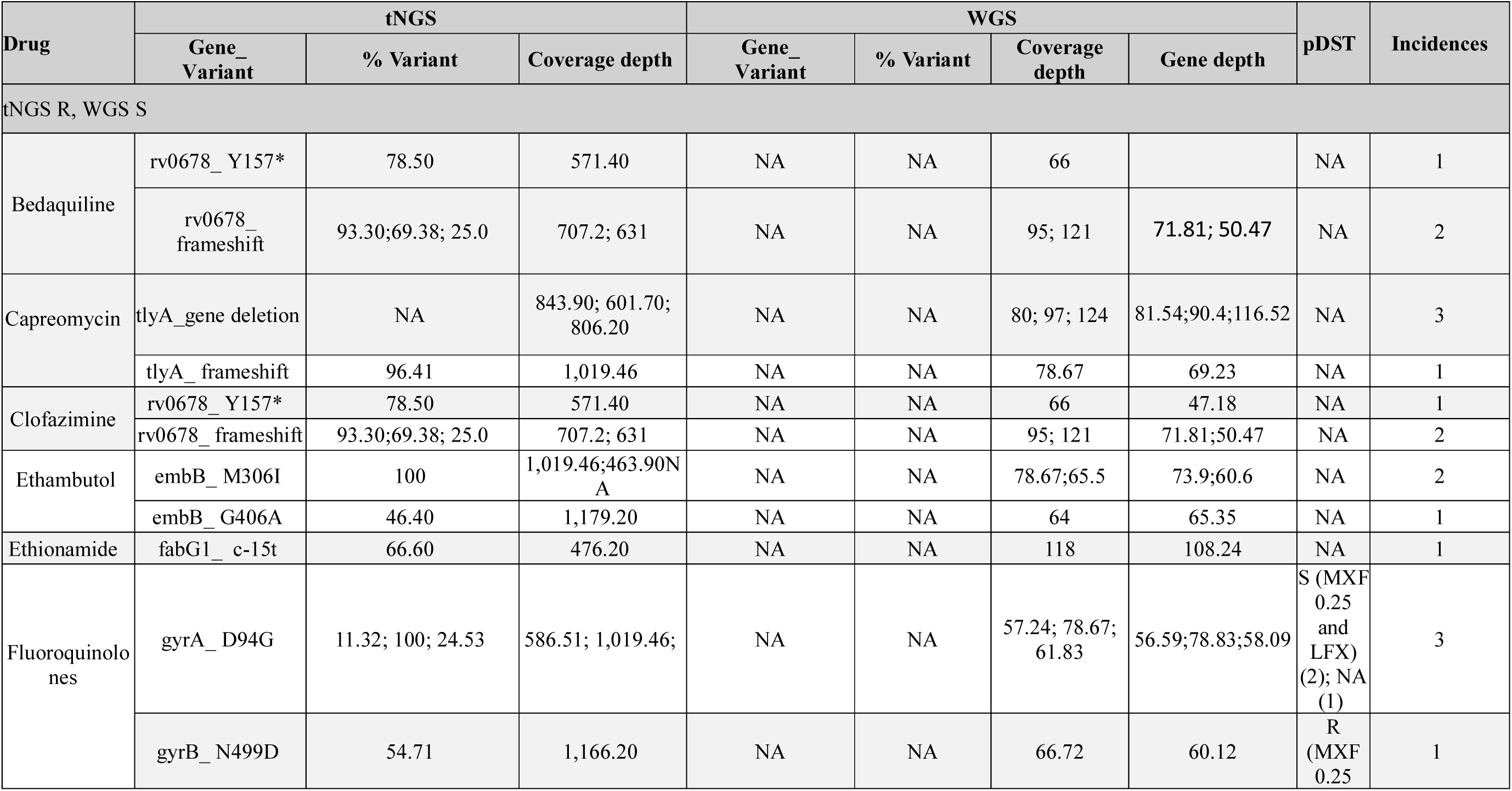
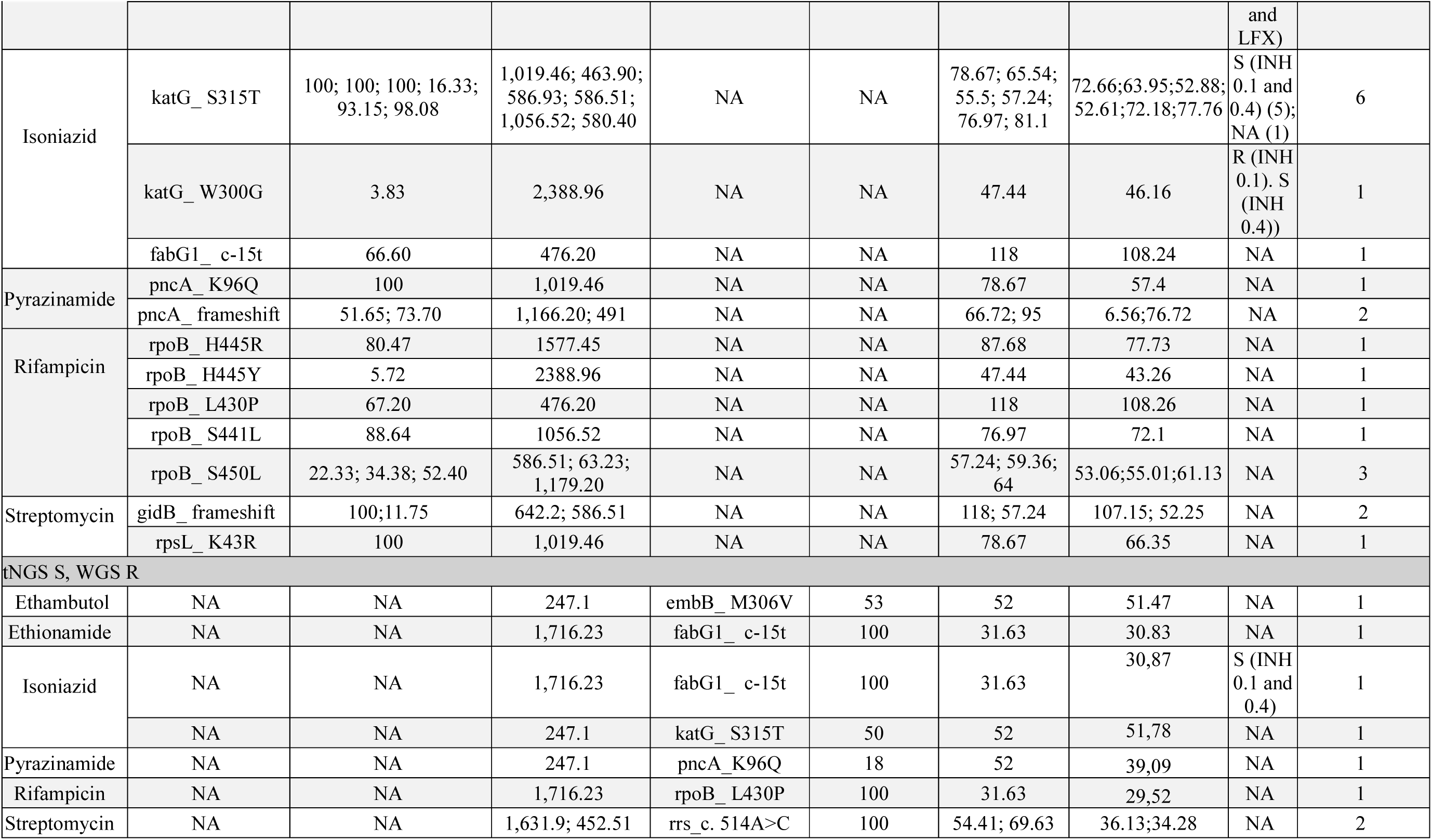
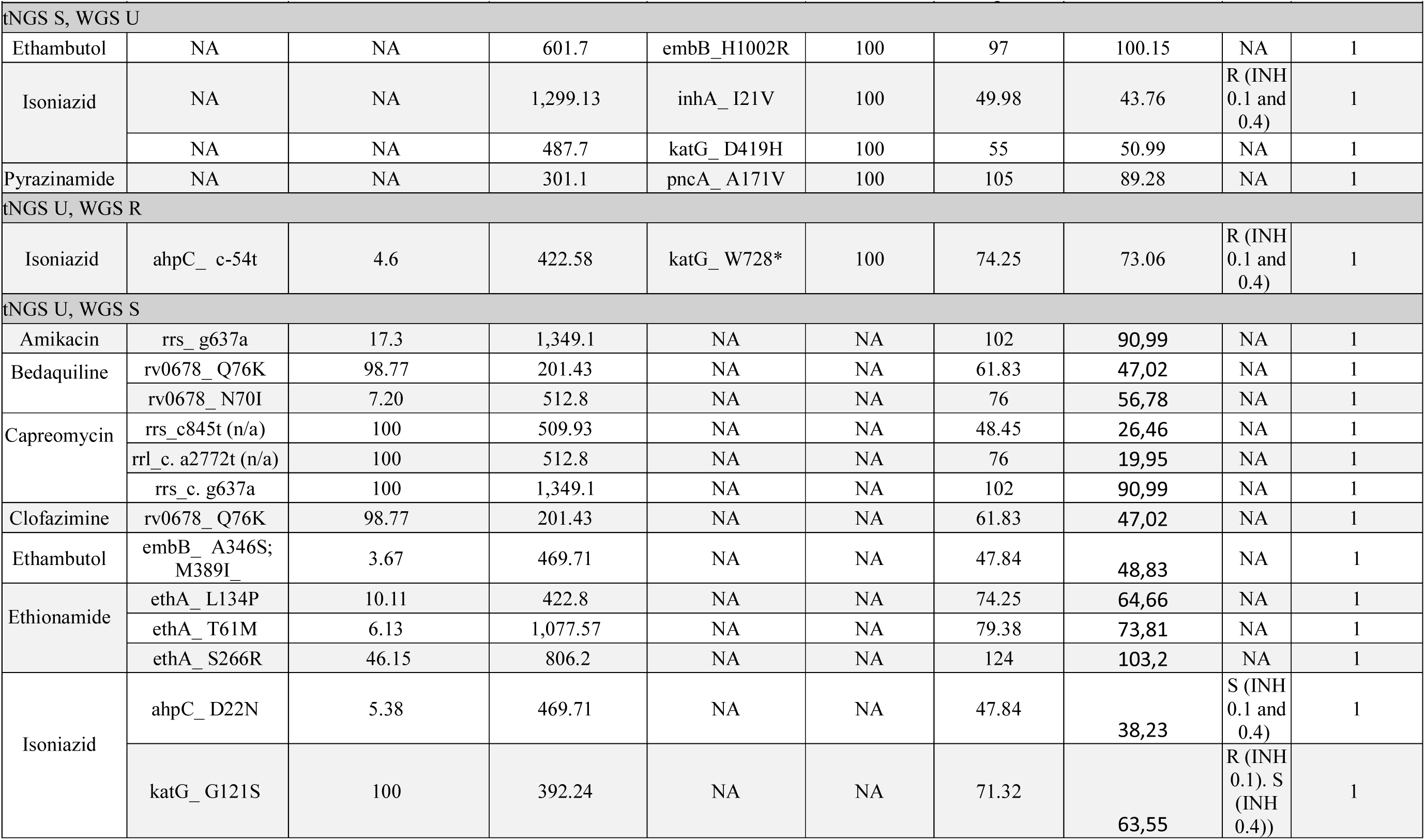
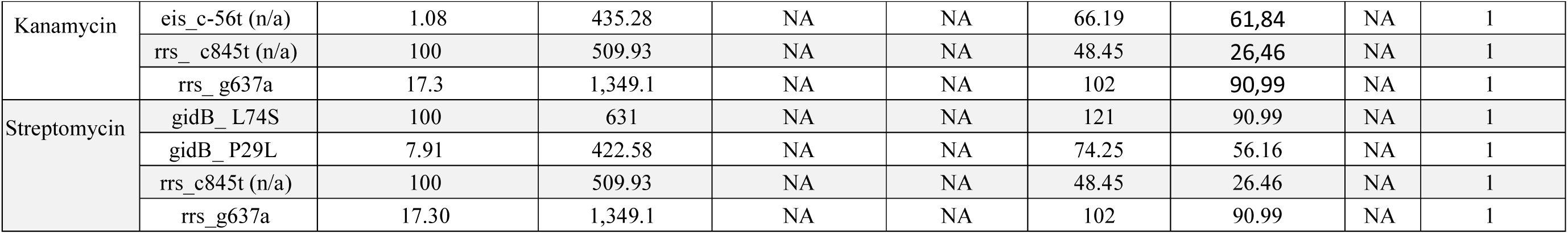
Table of discrepancies of WGS vs. tNGS results among 116 samples. Discrepancies were found affecting twelve anti-TB drugs. In the table below “S” stands for susceptible. and “R” stands for resistant. and “U” for Resistance – Uncertain Significant. “NA” means not available. In WGS column. it means no variant was detected that corresponds to the resistance status of interest. In pDST column. it means that pDST results were not available.

**Supplementary Material 2.**
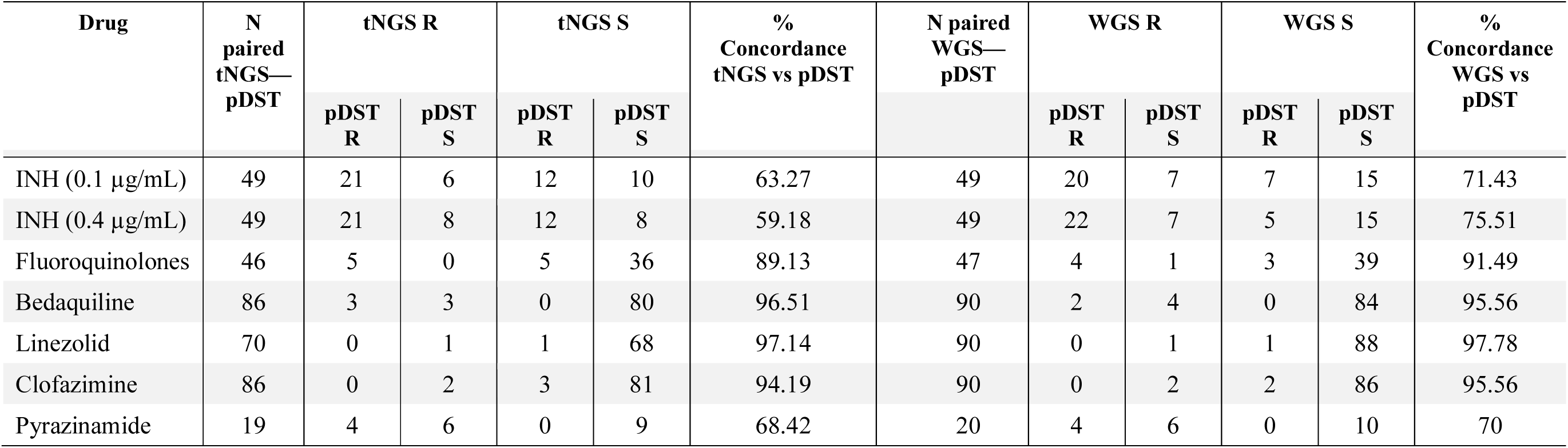
Table of detailed concordance between samples with available tNGS and pDST, and separately between samples with available WGS and pDST.

| Drug | N<br>paired<br>tNGS—<br>pDST | tNGS R |  | tNGS S |  | %<br>Concordance<br>tNGS vs pDST | N paired<br>WGS—<br>pDST | WGS R |  | WGS S |  | %<br>Concordance<br>WGS vs<br>pDST |
| --- | --- | --- | --- | --- | --- | --- | --- | --- | --- | --- | --- | --- |
|  |  | pDST<br>R | pDST<br>S | pDST<br>R | pDST<br>S |  |  | pDST<br>R | pDST<br>S | pDST<br>R | pDST<br>S |  |
| INH (0.1 µg/mL) | 49 | 21 | 6 | 12 | 10 | 63.27 | 49 | 20 | 7 | 7 | 15 | 71.43 |
| INH (0.4 µg/mL) | 49 | 21 | 8 | 12 | 8 | 59.18 | 49 | 22 | 7 | 5 | 15 | 75.51 |
| Fluoroquinolones | 46 | 5 | 0 | 5 | 36 | 89.13 | 47 | 4 | 1 | 3 | 39 | 91.49 |
| Bedaquiline | 86 | 3 | 3 | 0 | 80 | 96.51 | 90 | 2 | 4 | 0 | 84 | 95.56 |
| Linezolid | 70 | 0 | 1 | 1 | 68 | 97.14 | 90 | 0 | 1 | 1 | 88 | 97.78 |
| Clofazimine | 86 | 0 | 2 | 3 | 81 | 94.19 | 90 | 0 | 2 | 2 | 86 | 95.56 |
| Pyrazinamide | 19 | 4 | 6 | 0 | 9 | 68.42 | 20 | 4 | 6 | 0 | 10 | 70 |

**Supplementary Material 3.** Table of detailed information of available pDST results among samples with discrepant results (drug-level).

| No. | Drug | tNGS Result | WGS Result | pDST Results | Category |
| --- | --- | --- | --- | --- | --- |
| 1. | Isoniazid | U | S | S (INH 0.1 and 0.4) | Concordant with WGS |
| 2. | Isoniazid | S | R | S (INH 0.1 and 0.4) | Concordant with tNGS |
| 3. | Fluoroquinolones | R | S | S (MFX 0.25 and LFX) | Concordant with WGS |
| 4. | Fluoroquinolones | R | S | S (MFX 0.25 and LFX) | Concordant with WGS |
| 5. | Fluoroquinolones | R | S | R (MFX 0.25 and LFX) | Concordant with tNGS |
| 6. | Isoniazid | S | U | R (INH 0.1 and 0.4) | More likely to be concordant with WGS (as WGS still detected a resistance-associated mutation) |
| 7. | Isoniazid | R | S | S (INH 0.1 and 0.4) | Concordant with WGS |
| 8. | Isoniazid | R | S | S (INH 0.1 and 0.4) | Concordant with WGS |
| 9. | Isoniazid | R | S | S (INH 0.1 and 0.4) | Concordant with WGS |
| 10. | Isoniazid | R | S | S (INH 0.1 and 0.4) | Concordant with WGS |
| 11. | Isoniazid | R | S | S (INH 0.1 and 0.4) | Concordant with WGS |
| 12. | Isoniazid | R | S | R (INH 0.1), S (INH 0.4)) | Concordant with tNGS |
| 13. | Isoniazid | U | S | R (INH 0.1), S (INH 0.4)) | More likely to be concordant with tNGS (as tNGS still detected a resistance-associated mutation) |
| 14. | Bedaquiline | R | S | R (BDQ) | Concordant with tNGS |
| 15. | Clofazimine | R | S | S (CFZ) | Concordant with WGS |
| 16. | Isoniazid | U | R | R (INH 0.1 and 0.4) | Concordant with WGS |

